# A comparative study on promoter DNA methylation status of the *CDO1* tumor suppressor gene in chronic gastritis and gastric cancer: The role of *Helicobacter pylori* infection

**DOI:** 10.1101/2025.10.19.683320

**Authors:** Selin Kankaya, Metehan Karatas, Engin Hatipoglu, Nuray Kepil, Zulal Kaptan, Zeynep Caliskan, Matem Tuncdemir, Yildiz Dincer

## Abstract

Silencing of the *Cysteine dioxygenase-1 (CDO1*) tumor suppressor gene by aberrant DNA methylation contributes to gastric carcinogenesis. Despite earlier data indicating that *Helicobacter pylori* (*H. pylori*) infection induces aberrant DNA methylation in the gastric mucosa, the CDO1 methylation level has not been studied in *H. pylori*-positive (HPP) patients with chronic gastritis. This study aimed to investigate the *CDO1* methylation level in patients with chronic gastritis who have and do not have *H. pylori* infection, in comparison to gastric tumours.

The quantitative analysis of the promoter methylation of the *CDO1* gene and *CDO1* immunopositivity was determined in 45 primary gastric tumors, 17 biopsy samples of HPP- chronic gastritis, 23 *H. pylori*-negative (HPN) chronic gastritis, and 15 normal mucosa as the control. *CDO1* expression levels were measured by immunohistochemical staining, and the methylation level of the promoter region was determined by quantitative methylation-specific PCR after bisulfite conversion. *CDO1* gene promoter methylation levels were significantly higher in the gastric cancer (GC) group compared to the HPN-chronic gastritis and control groups (P = 0.000). There was no significant difference between the GC group and the HPP- chronic gastritis group. The *CDO1* gene promoter methylation levels were higher in the HPP- chronic gastritis group than in the HPN-chronic gastritis group (P = 0.049). There was no significant difference between the study groups for the *CDO1* immunopositivity score. The *CDO1* gene promoter methylation levels are increased in GC and HPP-chronic gastritis patients. The *CDO1* methylation, regardless of differences in protein expression, may be a promising non-invasive marker for early diagnosis of GC in HPP-chronic gastritis patients.

## Introduction

Chronic gastritis, particularly that associated with *Helicobacter pylori* (*H. pylori*) infection, is a critical premalignant stage in the multistep cascade leading to gastric cancer (GC) [1]. Genetic and epigenetic alterations play a crucial role in the development of GC. Among the key epigenetic mechanisms, DNA methylation is particularly significant in regulating gene expression. This process involves the addition of a methyl group to the carbon-5 position of the cytosine ring preceding a guanine nucleotide (CpG dinucleotide), resulting in the formation of 5-methylcytosine (5-CH₃-cytosine), and is catalysed by *DNA methyltransferases* (DNMTs). In gastric cancers, aberrant DNA methylation events are observed more frequently than mutations and other genetic abnormalities, highlighting the pivotal role of epigenetic modifications in the development and progression of the disease [2]. Since DNA methylation is a reversible epigenetic modification, targeting and restoring these abnormal methylation patterns represents a promising therapeutic strategy in cancer management.

Chronic inflammation induced by *H. pylori* infection promotes aberrant DNA methylation in gastric mucosal cells. Notably, hypermethylation of promoter CpG islands in tumor suppressor genes such as *SFRP1* [3], *CDKN2A* [4], *CDH1* [5], and *CDO1* [6] leads to their transcriptional silencing, thereby contributing to the development of GC.

Likely, epigenetic silencing of the mentioned tumor suppressor genes results from dysregulation of methylation-related genes due to exposure to cytokines produced by innate immunity [7–9]. The *TET* (*Ten-eleven translocation*) genes are involved in DNA demethylation. Activation of *NF-Κβ* pathway causes an increase in expression of *TET*-targeting miRNAs, which in turn downregulates *TET* expression. In addition, activities of DNMTs are increased by exposure to nitric oxide (NO) [8]. *NF-Κβ* activation and increased NO generation are consistently observed in H. *pylori*-induced gastritis [10].

*Cysteine dioxygenase 1* (*CDO1)* is a tumor suppressor gene. It encodes a non-heme, iron-dependent enzyme that regulates redox homeostasis through glutathione and promotes cell death under oxidative stress [11]. *CDO1* expression is epigenetically suppressed in tumor cells through promoter hypermethylation, which contributes to the evasion of apoptosis [12].

Hypermethylation in the promoter region of *CDO1* was determined in GC [6], but the methylation status of *CDO1* has not been examined in chronic gastritis yet. Given that gastric carcinoma often arises in the context of chronic gastritis and *H. pylori* positivity is frequently detected in the chronic gastritis, we hypothesised that aberrant methylation in the promoter region of *CDO1* may be a possible link between chronic gastritis and GC. In the present study, we examined the methylation level of the *CDO1* promoter region and the *CDO1* immunopositivity in primary gastric tumors and biopsy samples of chronic gastritis, which have and do not have *H. pylori* infection.

## Materials and Methods

### Study design and clinical samples

This study was conducted in accordance with the Declaration of Helsinki, and the protocol used was approved by the Clinical Research Ethics Committee of Cerrahpasa Medical Faculty (07/2020-83045809-604.01.02). All tissue samples were obtained from patients treated at Cerrahpaşa Medical Faculty Hospital between 10/11/2010, and 01/03/2020. All participants gave written informed consent before inclusion in the study.

The required sample size was calculated using G*Power 3.1.9.6 software. A minimum of 15 participants per study group was determined based on a one-way ANOVA, assuming a large effect size (f = 0.40), a significance level (α) of 0.05, and a power (1 – β) of 0.90. Accordingly, four study groups were formed, each consisting at least 15 cases: the control group (n=15), the *H. pylori*-negative (HPN)-chronic gastritis group (n=23), the *H. pylori*-positive (HPP)-chronic gastritis group (n=17), and the GC group (n=45).

The chronic gastritis groups, which include patients with and without *H. pylori* infection, are constituted by patients who undergo stomach endoscopic examination and are diagnosed with chronic gastritis based on pathological evaluation of biopsy samples. None of the chronic gastric patients has a malignant neoplasm. The control group is constituted by normal mucosa samples of the cases without *H. pylori* infection. The GC group consisted of patients diagnosed with primary gastric adenocarcinoma who underwent curative gastrectomy at Cerrahpasa Medical Faculty Hospital. Chemotherapy/radiotherapy received cases before the surgical procedure were excluded from the study. Formalin-fixed, paraffin-embedded (FFPE) samples obtained from the antrum, archived in the pathology department between 10/11/2010 and 01/03/2020, were used for methylation and immunohistochemical analysis. Tumor purity, the proportion of cancer cells in the tumor tissue, was evaluated by visual examination under a microscope by an expert pathologist. Each tissue has at least 30% tumor cells. The clinical classification of gastric carcinoma was performed according to the 2017 American Joint Committee on Cancer (AJCC) staging [13].

### Analysis of the *CDO1* gene promoter DNA methylation

%DNA methylation levels of the *CDO1* gene in primary gastric tumors and biopsy samples were determined by quantitative methylation-specific PCR (Q-MSP) after bisulfite conversion [14]. Genomic DNA extraction, bisulfite treatment, and Q-MSP were performed using commercial kits.

#### Genomic DNA extraction and bisulfite treatment

FFPE tissue was cut into six slices 5 μm thick, and genomic DNA extraction was performed using the Quick-DNA Miniprep Plus Kit according to the FFPE tissue protocol (ZymoResearch, Irvine, CA, USA, Cat no: D4068). Following the protocol, after FFPE tissues were deparaffinized, DNA was extracted using proteinase K. The concentration and purity values of the isolated DNA samples were measured with the NanoDrop ScientificTM 1000 (Thermo Fisher, Delaware, USA) device.

Bisulfite modification was performed on DNA samples (1 µg) isolated from paraffin blocks using the EZ DNA Methylation-GoldTM Kit (Zimo Research, Irvine, CA, USA, Cat no: D5005). In principle, while all unmethylated cytosines in DNA are converted to uracil by chemical reaction, methylated cytosines in DNA remain unchanged. After this conversion process was completed, the DNA was amplified by PCR. During PCR amplification, thymine is placed where the uracils are located in the DNA sequence, and methylated cytosines remain in the form of cytosines. Since unmethylated cytosine is converted to thymine, all cytosines read in the DNA sequence represent methylation sites. The methylation profiles were determined using forward and reverse methylated primers specifically designed for methylation sites on the *CDO1* promoter. Primers were designed to be specific to the *CDO1* promoter region, ensuring that they contained sequences that would bind only to the target region to prevent potential non-specific binding. In addition to the recommendations of the design tools used, the regions affected by the primers were examined in detail in FASTA format, and the results were validated with relevant NCBI tools.

#### Determination of the amount of DNA required for methylation analysis and preparation of methylation standards

The optimal amount of DNA per bisulfite treatment is 200 to 500 ng according to the commercial kit protocol. 200 ng of DNA was used in the study. The required amount of DNA samples was pipetted into PCR microplate wells. The final volume was completed with ultrapure water to 20 μl. Commercial 100% Methylated and non-methylated human DNA standards (Zymo Research, Irvine, CA, USA. Cat. no: D5014) were converted with bisulfite as controls of methylation-specific PCR assays. The standards were prepared from bisulfite-converted methylated and unmethylated DNA sets containing percentages of fully methylated DNA (100, 75, 50, 25, 10, 5, and 0). The standard curve chart was created with the Curve Expert program according to the methylation percentages determined according to the cycle threshold (Ct) values of the standards.

#### Quantitative methylation-specific PCR (Q-MSP)

We performed real-time PCR using iQ Supermix (Bio-Rad, Hercules, CA) real-time PCR System and real-time PCR Thermal Cycler (Roche Diagnostics) for dedection of *CDO1* promoter methylation levels. After bisulfite conversion, DNA was amplified by real-time PCR using methylated forward (5’-TTTGGGACGTCGGAGATAAC-3’) and reverse primers (5’- CCAACATTAAAATACCGAAACGTA-3’). The Q-MSP reaction (20 µl / well) contained 10 µl of Zymo TaqTM qPCR PreMix (Zymo Research, Irvine, CA, USA. Cat. no: E2055) that includes a hot-start DNA polymerase and buffer system optimised for the amplification of bisulfite-treated DNA and an intense double-strand DNA-specific fluorescent dye, SYTO 9®, for sensitive real-time DNA quantification, 2.5 µl assay-specific forward and reverse primers for the methylated loci, and bisulfite-treated DNA, as a template and labelled probs for *CDO1* unmethylated and *CDO1* methylated. The PCR reactions were incubated at 95°C for 10 min, followed by 40 cycles at 95°C for 15 sec and 55°C for 1 min. PCRs were performed in duplicate. At the end of the reaction quantitative analysis of methylated alleles was defined as previously described [15]. The Ct values were converted to per cent methylation using the standard curve chart.

#### Immunostaining for *CDO1* protein in tissue samples

FFPE tissue blocks were cut into 5 μm thick sections, and immunohistochemistry was performed as described previously [16]. The sections were incubated with anti-*CDO1* polyclonal antibody [CUSABIO CSB-PA438445 Cysteine Dioxygenase, Type I [Polyclonal] Conc. 0.1 ml (1 : 25 – 100)]. The secondary antibody reaction was performed using the Histostain-Plus Bulk Kit (Invitrogen, Camarillo, USA). Immunoreactive products were visualised by 3,3-diaminobenzidine catalysis. *CDO1* protein levels were evaluated using the immunoreactive-score (IRS) system in primary gastric tumors and biopsy samples of chronic gastritis [17]. Cells were scored by assessment of the number of positive cells and the percentage of staining. The *CDO1* immunoreactivity was scored based on percentages of the stained area. Then the percentage of positive cells was scored as follows: 0 (negative, no staining measured), 1 (low [1-40% staining area]), 2 (medium [40-60% staining area]), 3 (high [60-80% staining area]), 4 (very high [80-100% staining area]). Positively stained cells were counted under ×20 magnification in 12 randomly selected fields, and the immunoreactivity scores of each were determined and averaged to obtain a single numerical value. Representative immunostaining of the *CDO1* protein in gastric tumors is presented in Fig 1.

**Fig 1.**
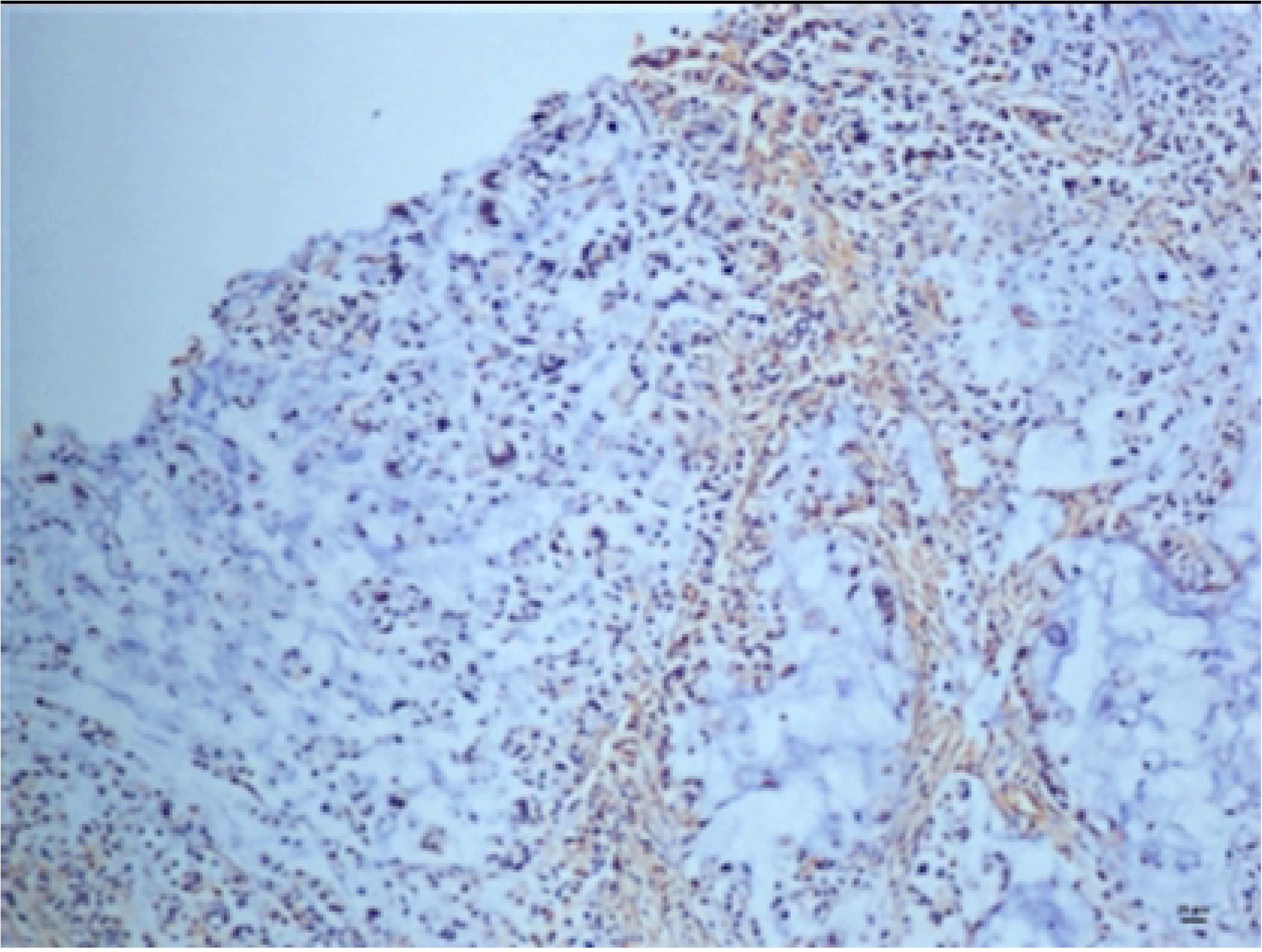
Representative immunostaining of the CDO1 in the gastric tumors. (A) Low immunoreaction (B) Medium immunoreaction (C) High immunoreaction (D) Negative control staining (E) Positive control staining

### Statistical analyses

The data were analyzed using SPSS version 22.0 for Windows. The Shapiro-Wilk test was used to determine the distribution of the data. *CDO1* methylation levels and immunopositivity were not normally distributed, therefore analyzes were performed by the nonparametric Kruskal-Wallis and Mann-Whitney U tests. Spearman correlation analysis evaluated the relationship between dependent variables, as appropriate. The results were presented as the median (min-max). The optimal cut-off value for % *CDO1* methylation in predicting GC was determined through receiver operating characteristic (ROC) curve analysis, and the area under the curve (AUC) was estimated. The cut-off value with the highest accuracy was determined through a plot of sensitivity/specificity against the criterion value. P value < 0.05 was considered statistically significant.

## Results

The demographic data of the study groups and the characteristics of FFPE samples are presented in Tables 1 and 2, respectively. No significant gender difference was found for *CDO1* gene promoter methylation levels and immunopositivity scores in the study groups (Table 1). There was no significant correlation between age and *CDO1* gene promoter methylation levels and immunopositivity scores in the study groups.

**Table 1.** The demographic data of the study groups.

|  | Control (n=15) | HPN (n=23) | HPP (n=17) | GC (n=45) | P value |
| --- | --- | --- | --- | --- | --- |
| **Sex, n (%)** |  |  |  |  |  |
| **Female** | 6 (40.0) | 12 (52.1) | 9 (52.9) | 17 (37.7) | 0.72 |
| **Male** | 9 (60.0) | 11 (47.8) | 8 (47.1) | 28 (62.3) |  |
| **Age, median (IQR)** | 65 (46.0-77.0) | 64 (46.0-77.0) | 59 (36.0-84.0) | 61 (35.0-86.0) | 1.00 |

Control: Healthy people without *H. pylori* infection, HPN: *H. pylori*-negative chronic gastritis, HPP: *H. pylori*-positive chronic gastritis, GC: Gastric cancer. IQR: interquartile ranges, P values correspond to the Kruskal-Wallis test (depending on distribution) and statistically significant differences are P < 0.05.

**Table 2.** Characteristics of FFPE samples used for DNA methylation and immunohistochemical analyses.

| Tissue | Case no | Age | Gender | Cancer | TNM clasification | Stage |
| --- | --- | --- | --- | --- | --- | --- |
| **Normal** | Normal-1 | 71 | F | - | - | - |
|  | Normal-2 | 51 | M | - | - | - |
|  | Normal-3 | 61 | M | - | - | - |
|  | Normal-4 | 57 | F | - | - | - |
|  | Normal-5 | 76 | M | - | - | - |
|  | Normal-6 | 77 | M | - | - | - |
|  | Normal-7 | 58 | M | - | - | - |
|  | Normal-8 | 69 | M | - | - | - |
|  | Normal-9 | 64 | F | - | - | - |
|  | Normal-10 | 68 | M | - | - | - |
|  | Normal-11 | 46 | F | - | - | - |
|  | Normal-12 | 65 | F | - | - | - |
|  | Normal-13 | 76 | M | - | - | - |
|  | Normal-14 | 74 | M | - | - | - |
|  | Normal-15 | 53 | F | - | - | - |
| **HPN** | HPN-1 | 71 | F | - | - | - |
|  | HPN-2 | 51 | M | - | - | - |
|  | HPN-3 | 59 | F | - | - | - |
|  | HPN-4 | 70 | F | - | - | - |
|  | HPN-5 | 61 | M | - | - | - |
|  | HPN-6 | 64 | F | - | - | - |
|  | HPN-7 | 57 | F | - | - | - |
|  | HPN-8 | 63 | F | - | - | - |
|  | HPN-9 | 76 | M | - | - | - |
|  | HPN-10 | 64 | F | - | - | - |
|  | HPN-11 | 71 | M | - | - | - |
|  | HPN-12 | 77 | M | - | - | - |
|  | HPN-13 | 58 | M | - | - | - |
|  | HPN-14 | 69 | M | - | - | - |
|  | HPN-15 | 64 | F | - | - | - |
|  | HPN-16 | 68 | M | - | - | - |
|  | HPN-17 | 46 | F | - | - | - |
|  | HPN-18 | 65 | F | - | - | - |
|  | HPN-19 | 62 | M | - | - | - |
|  | HPN-20 | 76 | M | - | - | - |
|  | HPN-21 | 74 | M | - | - | - |
|  | HPN-22 | 63 | F | - | - | - |
|  | HPN-23 | 53 | F | - | - | - |
| **HPP** | HPP-1 | 60 | M | - | - | - |
|  | HPP-2 | 84 | F | - | - | - |
|  | HPP-3 | 59 | M | - | - | - |
|  | HPP-4 | 51 | M | - | - | - |
|  | HPP-5 | 55 | M | - | - | - |
|  | HPP-6 | 64 | F | - | - | - |
|  | HPP-7 | 71 | M | - | - | - |
|  | HPP-8 | 51 | F | - | - | - |
|  | HPP-9 | 75 | F | - | - | - |
|  | HPP-10 | 47 | F | - | - | - |
|  | HPP-11 | 47 | M | - | - | - |
|  | HPP-12 | 36 | F | - | - | - |
|  | HPP-13 | 59 | F | - | - | - |
|  | HPP-14 | 58 | F | - | - | - |
|  | HPP-15 | 78 | M | - | - | - |
|  | HPP-16 | 52 | F | - | - | - |
|  | HPP-17 | 65 | M | - | - | - |
| **GC** | Cancer-1 | 57 | M | + | T4AN2 | IIIB |
|  | Cancer-2 | 62 | M | + | T4AN0 | IIB |
|  | Cancer-3 | 55 | M | + | T3N1 | IIB |
|  | Cancer-4 | 69 | F | + | T1AN0 | IA |
|  | Cancer-5 | 55 | M | + | T2N2 | IIB |
|  | Cancer-6 | 79 | M | + | T1BN0 | IA |
|  | Cancer-7 | 69 | M | + | T4N3B | IIIC |
|  | Cancer-8 | 48 | M | + | T4N3 | IIIC |
|  | Cancer-9 | 50 | F | + | T4AN3A | IIIC |
|  | Cancer-10 | 81 | F | + | T3N1 | IIB |
|  | Cancer-11 | 58 | F | + | T2N0 | IB |
|  | Cancer-12 | 56 | F | + | T4AN3 | IIIC |
|  | Cancer-13 | 47 | M | + | T2N2 | IIB |
|  | Cancer-14 | 39 | F | + | T2N0 | IB |
|  | Cancer-15 | 35 | M | + | T1N0 | IA |
|  | Cancer-16 | 71 | M | + | T4AN0 | IIB |
|  | Cancer-17 | 45 | F | + | T4AN3 | IIB |
|  | Cancer-18 | 52 | F | + | T4AN3 | IIIC |
|  | Cancer-19 | 59 | M | + | T1N0 | IA |
|  | Cancer-20 | 50 | M | + | T4AN3 | IIIC |
|  | Cancer-21 | 50 | M | + | T4N0 | IIB |
|  | Cancer-22 | 61 | M | + | T3N3 | IIIB |
|  | Cancer-23 | 66 | F | + | T3N1 | IIB |
|  | Cancer-24 | 51 | F | + | T4N3 | IIIC |
|  | Cancer-25 | 76 | F | + | T3N1 | IIB |
|  | Cancer-26 | 63 | M | + | T2N2 | IIB |
|  | Cancer-27 | 67 | M | + | T4N3 | IIIC |
|  | Cancer-28 | 55 | M | + | T1N0 | IA |
|  | Cancer-29 | 65 | M | + | T1BN2 | IIA |
|  | Cancer-30 | 83 | F | + | T4AN0 | IIB |
|  | Cancer-31 | 48 | F | + | T4AN3 | IIIC |
|  | Cancer-32 | 86 | M | + | T4AN3 | IIIC |
|  | Cancer-33 | 82 | M | + | T4AN0 | IIB |
|  | Cancer-34 | 70 | M | + | T4AN3 | IIIC |
|  | Cancer-35 | 68 | M | + | T3N0 | IIA |
|  | Cancer-36 | 45 | M | + | T4AN0 | IIB |
|  | Cancer-37 | 57 | M | + | T4AN3 | IIIC |
|  | Cancer-38 | 70 | M | + | T4AN3 | IIIC |
|  | Cancer-39 | 71 | M | + | T4N3A | IIIC |
|  | Cancer-40 | 62 | M | + | T3N0 | IIA |
|  | Cancer-41 | 56 | M | + | T4AN3 | IIIC |
|  | Cancer-42 | 74 | F | + | T1AN0 | IA |
|  | Cancer-43 | 63 | F | + | T1AN0 | IA |
|  | Cancer-44 | 42 | F | + | T1BN2 | IIA |
|  | Cancer-45 | 61 | F | + | T4AN3 | IIIC |

F: Female, M: Male, Normal: Healthy people without *H. pylori* infection as control, HPN: *H. pylori*-negative chronic gastritis, HPP: *H. pylori*-positive chronic gastritis, GC: Gastric cancer. TNM: Tumor, Node, Metastasis.

The cytoplasmic *CDO1* immunopositivity was observed in all tissue samples. There was no significant difference between the study groups for the *CDO1* immunopositivity score (Table 3).

**Table 3.**
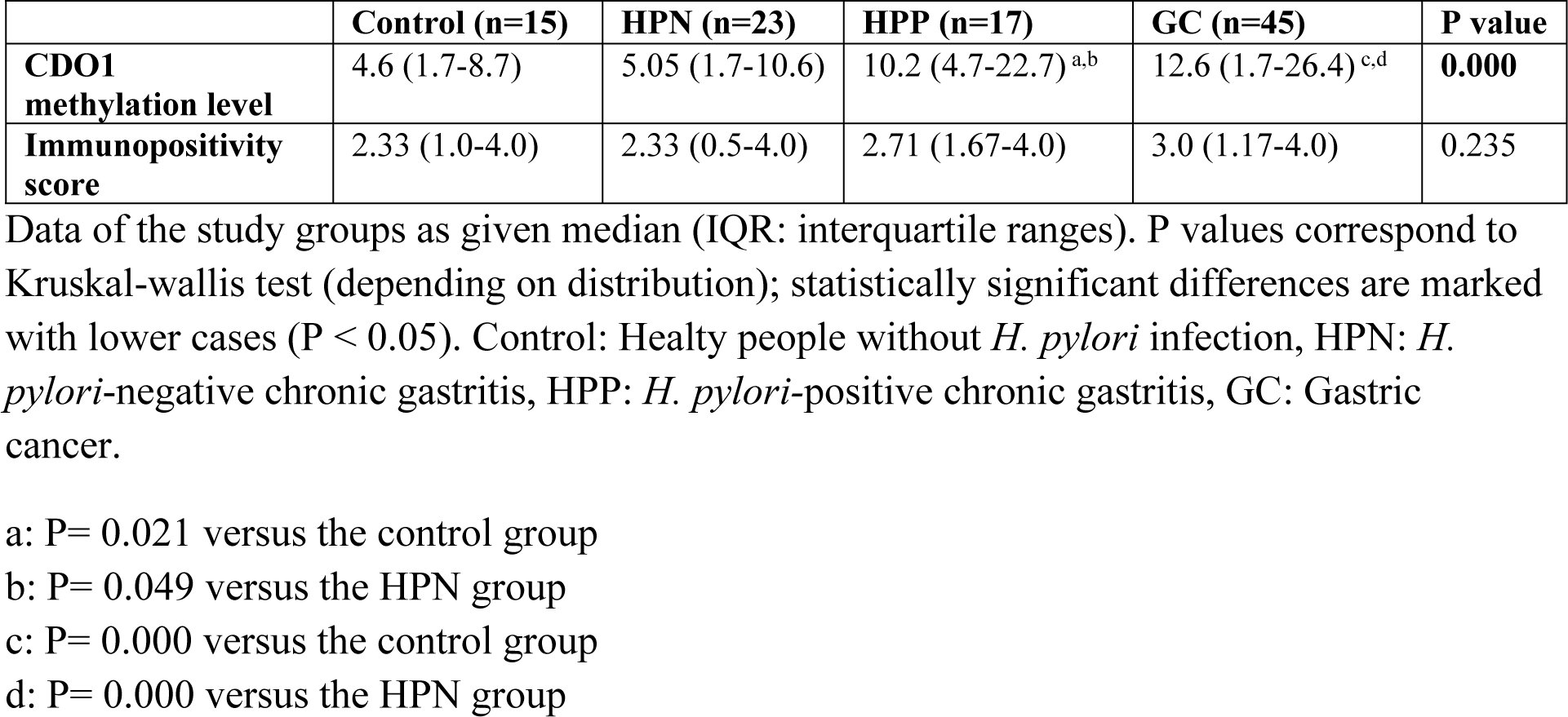
CDO1 promoter methylation levels and immunopositivity scores of the study groups.

Data of the study groups as given median (IQR: interquartile ranges). P values correspond to Kruskal-wallis test (depending on distribution); statistically significant differences are marked with lower cases (P < 0.05). Control: Healty people without *H. pylori* infection, HPN: *H. pylori*-negative chronic gastritis, HPP: *H. pylori-*positive chronic gastritis, GC: Gastric cancer.

*CDO1* gene promoter methylation levels were significantly higher in the GC group compared to the HPN-chronic gastritis and control groups (P = 0.000) (Table 3). There was no significant difference between the GC group and the HPP-chronic gastritis group. The *CDO1* gene promoter methylation levels were higher in the HPP-chronic gastritis group than in the HPN-chronic gastritis group (P = 0.049) (Table 3). There was no significant correlation between

*CDO1* gene promoter methylation levels and *CDO1* immunopositivity in the study groups. Distribution of the *CDO1* gene promoter methylation levels of the study groups is given in Fig 2.

**Fig 2.**
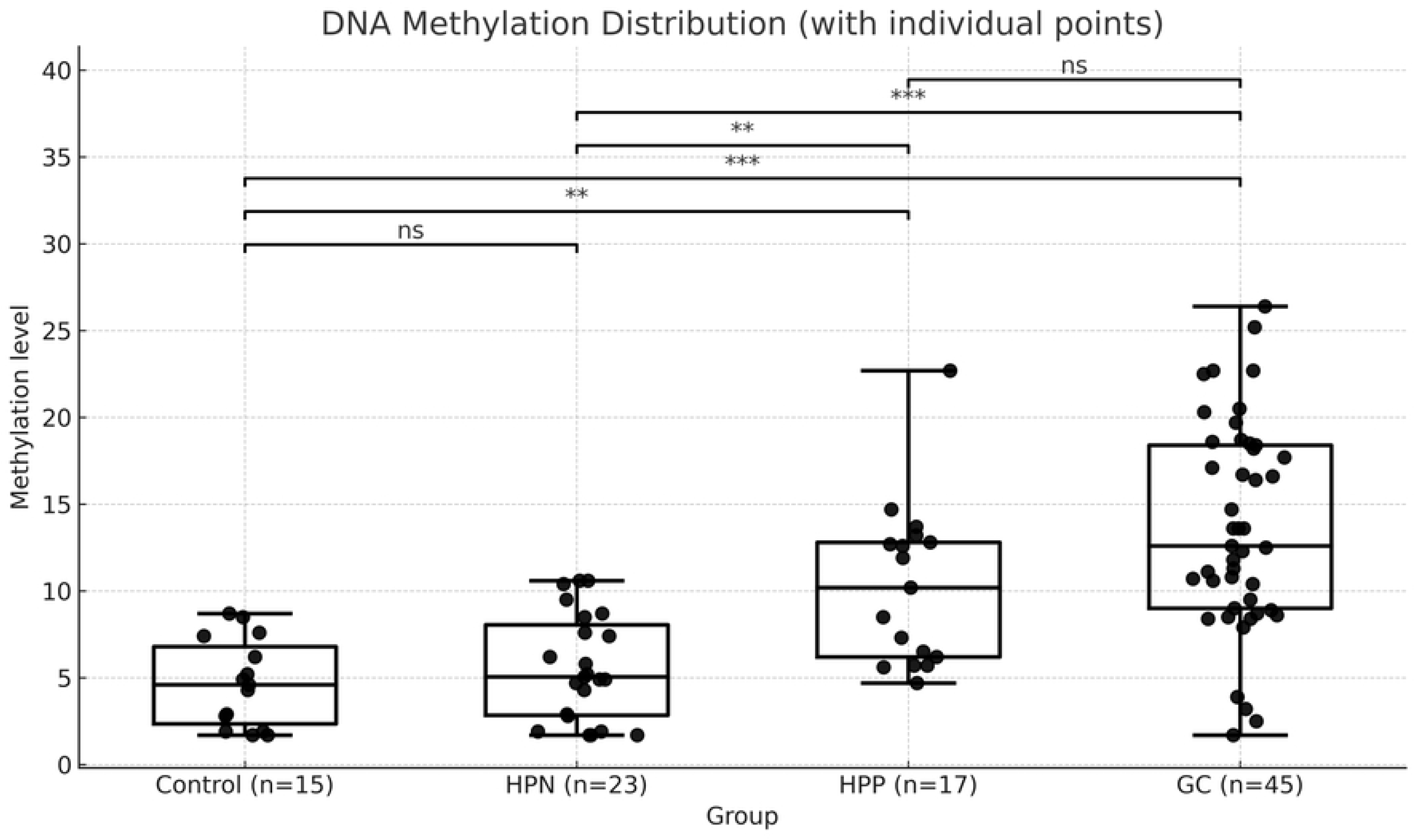
Distribution of the CDO1 gene promoter methylation levels of the study groups. Control: Healthy people without *H. pylori* infection, HPN: *H. pylori*-negative chronic gastritis, HPP: *H. pylori-*positive chronic gastritis, GC: Gastric cancer. P values correspond to Kruskal-Wallis and Dunn post-hoc test (depending on distribution); statistically significant differences are marked with stars. * p < 0.05 ** p < 0.01 *** p < 0.001 ns: not significant

The GC group was categorised into clinical stages as stage I, stage II, and stage III according to the 2017 AJCC staging [13]. The methylation levels of the *CDO1* gene promoter tended to be higher as the clinical stage progressed. There was a significant difference between the groups of stage III and stage I / II, respectively (P = 0.000, P=0.000) (Fig. 3).

**Fig 3.**
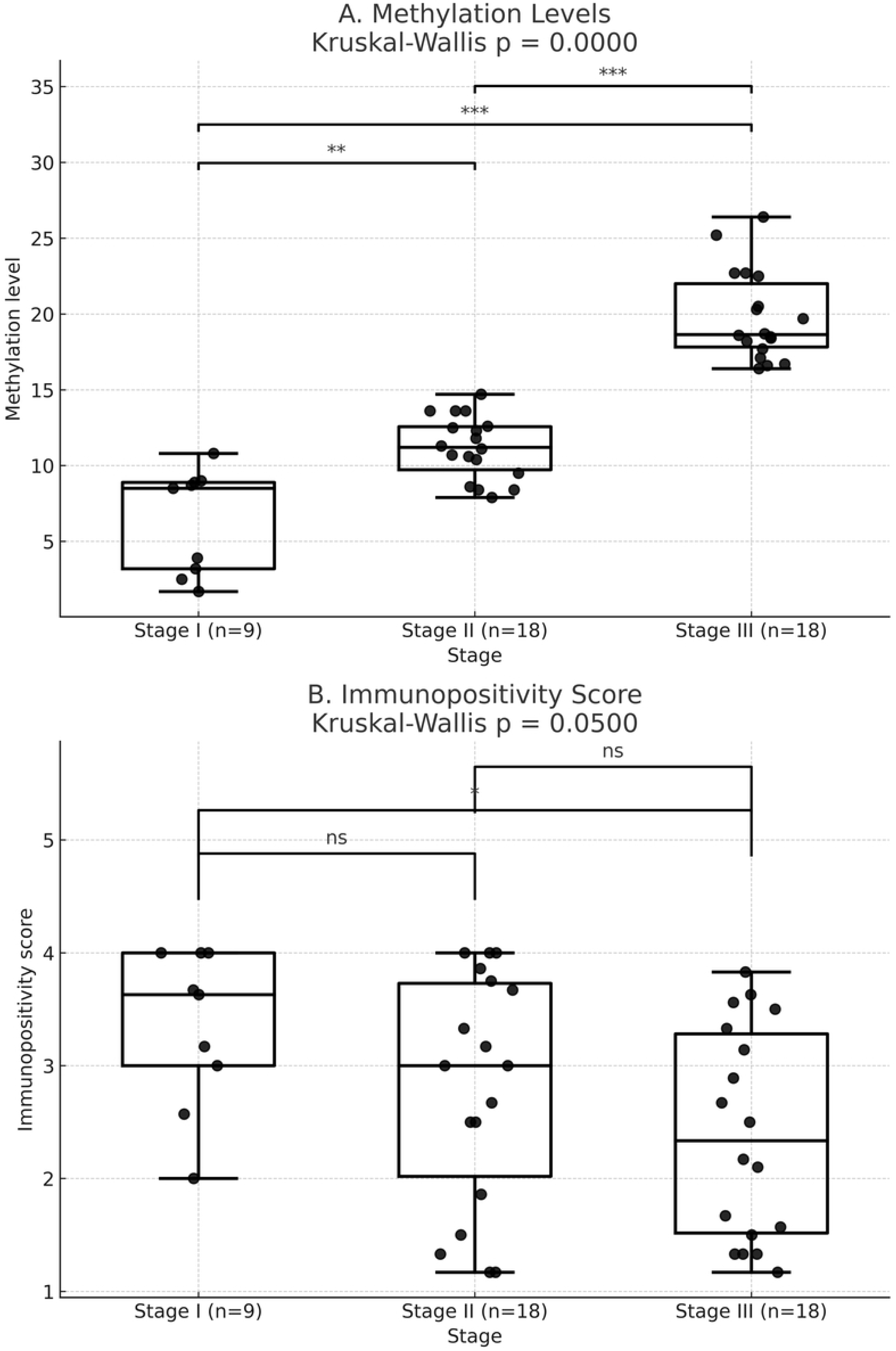
The CDO1 (A) promoter methylation levels and (B) immunopositivity scores according to the clinical stage of gastric cancer. P values correspond to Kruskal-Wallis and Dunn post-hoc test (depending on distribution); statistically significant differences are marked with stars. * p < 0.05 ** p < 0.01 *** p < 0.001 ns: not significant.

The ROC curve was used to assess the clinical performance of *CDO1* methylation. Two receiver operating characteristic (ROC) curves were drawn. First, a ROC curve was generated to evaluate the discriminative power of *CDO1* methylation levels between individuals with GC and those without cancer. The model demonstrated strong discriminative ability, with an AUC of 0.83, indicating that *CDO1* methylation levels could effectively differentiate between the two groups (95% CI: 0.74–0.91, p < 0.0001). This suggests that methylation levels have high diagnostic accuracy in distinguishing between cancer patients and non-cancer individuals. The optimal threshold was determined using the Youden Index, which identified a cut-off value of 7.90% DNA methylation (91.1 % sensitivity and 67.3% specificity). This suggests that individuals with DNA methylation levels above 7.90% are more likely to be classified as cancer cases with high sensitivity and moderate specificity (Fig. 4A). To assess the association between methylation levels and cancer status, a binary logistic regression analysis was conducted (odds ratio: 1.30, 95% CI: 1.16 – 1.44, P< 0.001). The results indicate that methylation is a statistically significant predictor of cancer. According to these data, for each one-unit increase in methylation level, the odds of having cancer increase by approximately 30%. This association is statistically significant (P < 0.001).

**Fig 4.**
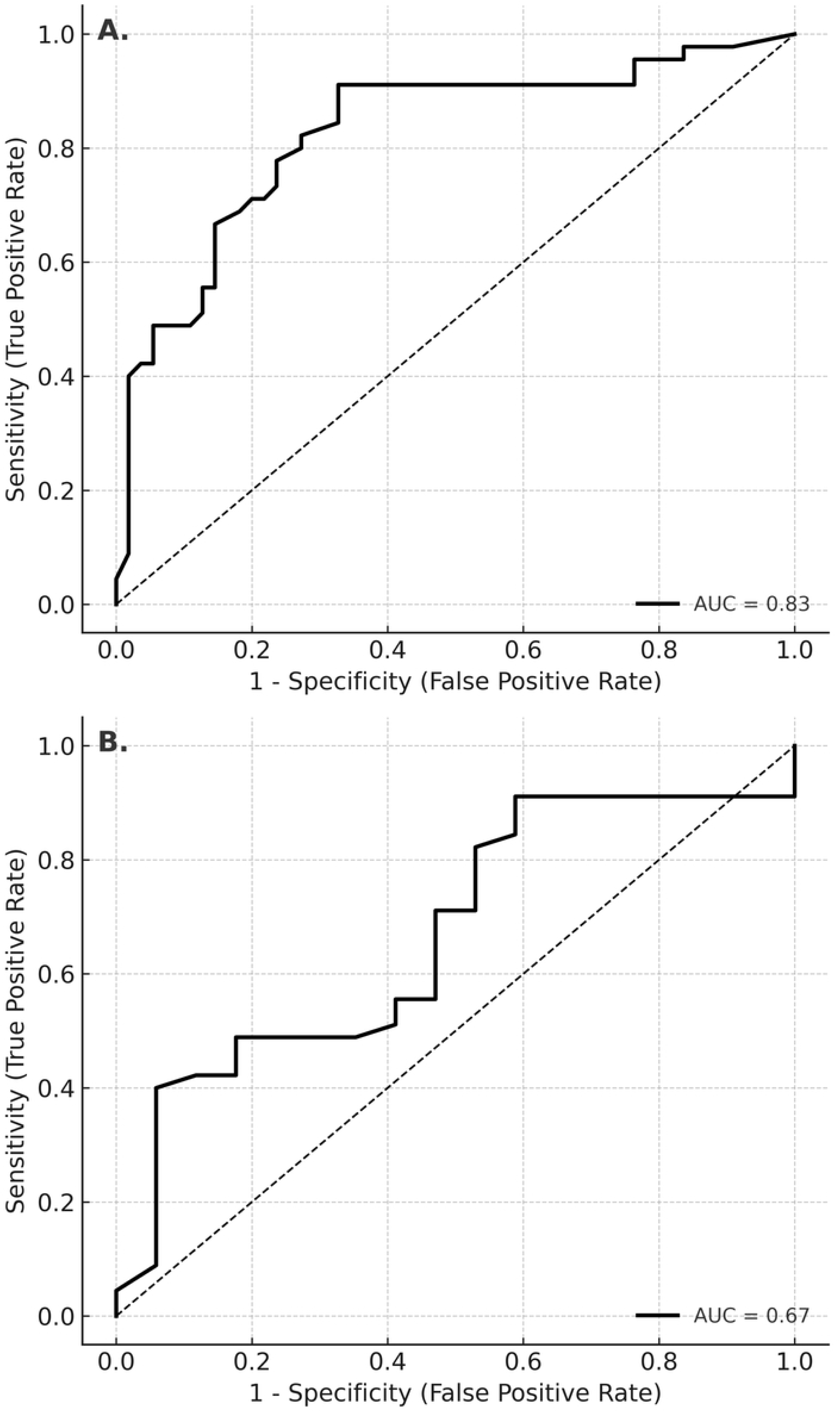
ROC analysis for (A) gastric cancer and non-cancer cases (B) gastric cancer and HPP-chronic gastritis cases. (A): AUC: 0.83, Cut-off: 7.90 % (91.1 % sensitivity and 67.3% specificity). (B): (A): AUC: 0.67, Cut-off: 7.90 % (91.1 % sensitivity and 41.2% specificity) or cut-off: 16.4 % (40.0 % sensitivity and 94.1 % specificity).

Second, a ROC curve was generated to assess the clinical performance of *CDO1* methylation in the differential diagnosis of GC and HPP-chronic gastritis (Fig 4B). The ROC analysis demonstrated an AUC of 0.67 (95% CI: 0.512–0.815, p = 0.011), indicating that the model’s discriminatory ability is statistically significant and moderate in strength. Based on the Youden index, the optimal cut-off value was identified as 7.90. At this threshold, the sensitivity and specificity of the test were calculated as 91.1% and 41.2%, respectively. These findings suggest that the test exhibits high sensitivity in detecting cancer cases, while its specificity in excluding HPP individuals remains limited. As an alternative to the cut-off value of 7.90, the Youden index identified 16.4 as another potential threshold. Based on ROC curve analysis, the model’s discriminatory ability was found to be moderate (AUC = 0.67; 95% CI: 0.512–0.815; p = 0.011). At the 16.4 threshold, the sensitivity and specificity of the test were calculated as 40.0% and 94.1%, respectively, suggesting higher specificity at the expense of reduced sensitivity.

## Discussion

In an early study, global DNA methylation of gastric mucosa was examined by immunohistochemical detection of 5-CH_3_ cytosine. The authors determined that global DNA methylation gradually decreased from normal mucosa to HPP chronic atrophic gastritis, there was a significant difference between HPN subjects and HPP-chronic atrophic gastritis. In the 10 patients with preneoplastic lesions, global DNA methylation decreased over time despite the *H. pylori* eradication, reaching significance at 10 years versus baseline [18]. In a recent study [19], methylation levels of some GC-related genes, *MOS*, miR124a-3, *NKX6-1*, *EMX1*, *CDH1*, and *TWIST1* in the non-cancerous gastric mucosa were compared between subjects with and without family history based on GC and *H. pylori* infection, and the changes in the methylation levels were evaluated over time after *H. pylori* eradication. They determined that the methylation of *MOS* and *CDH1* was associated with aberrant DNA methylation and gastric carcinogenesis in a family history of GC, and only *CDH1* methylation decreased after *H. pylori* eradication. As far as we know, there is no report about the *CDO1* methylation level in chronic gastritis with or without *H. pylori* infection. In this study, we revealed *H. pylori*-induced aberrant methylation in the *CDO1* promoter in gastric mucosa samples. *CDO1* gene promoter methylation levels were significantly higher in the HPP-chronic gastritis samples than in HPN- chronic gastritis samples. This is the first report that shows a DNA methylation change induced by *H. pylori* in the *CDO1* gene promoter, and indicates that an epigenetic mechanism is involved in chronic gastritis-induced GC.

Promoter DNA methylation of *CDO1* has gained great interest as a biomarker in GC. *CDO1* hypermethylation in peritoneal lavage fluid cells of stage IV GC patients was determined by Ushiku et al [20]. Kojima et al [6] reported that the promoter DNA methylation level of *CDO1* is predictive in detecting remnant GC after initial gastrectomy for primary GC. Kubota et al [21] suggested *CDO1* hypermethylation as a potential marker for metachronous GC, a recently developed GC emerging at a previously unaffected site at least one year after endoscopic submucosal dissection of the primary tumor. These previous studies have suggested the prognostic significance of *CDO1* promoter methylation in end-stage GC patients. However, there have been no reports on the *CDO1* promoter methylation level as a biomarker in the early detection of primary GC, especially in the follow-up of individuals with high-risk conditions such as chronic gastritis with *H. pylori* positivity. The information about the link between *H. pylori* positivity and *CDO1* promoter methylation status in chronic gastritis patients is insufficient. We are the first to investigate the promoter methylation levels of the *CDO1* gene in patients with GC and chronic gastritis with and without *H. pylori* infection, comparatively. As consistent with the results of the other studies [6, 20, 21], we found that *CDO1* promoter methylation levels in gastric tumors are significantly increased as the clinical stage, tumor size, and lymph node involvement progress. ROC analysis demonstrated that the *CDO1* methylation levels are both a highly accurate discriminator (AUC = 0.83) and a significant risk indicator for cancer (OR = 1.30, p < 0.001). The findings support the potential utility of *CDO1* methylation as a diagnostic biomarker in GC detection.

As the most important finding of the present study, we determined that *CDO1* promoter methylation levels are higher in primary gastric tumors than those in biopsy samples of chronic gastritis without *H. pylori* infection, whereas they are not significantly different from those in biopsy samples of chronic gastritis with *H. pylori.* According to ROC analysis, *CDO1* promoter methylation level exhibits high sensitivity (91.1%) in detecting cancer cases, while its specificity in excluding HPP individuals remains limited (41.2%) with the 7.90 cut-off value. This cut-off value may be more appropriate for screening purposes, where maximising sensitivity is prioritised, rather than for confirmatory diagnostic use, which requires higher specificity.

Alternatively, with the 16.4 cut-off value, the sensitivity and specificity of the test were calculated as 40.0% and 94.1%, respectively, suggesting higher specificity at the expense of reduced sensitivity. These findings indicate that the test has limited sensitivity for detecting cancer cases, but demonstrates high specificity in correctly excluding HPP individuals. The 16.4 cut-off value may be appropriate in settings where diagnostic precision is prioritised. Values equal to or above 16.4 could be defined as a ‘high-risk group,’ thereby contributing to clinical risk stratification. This classification may facilitate patient triage, determine follow-up frequency, and guide prioritisation. With a specificity of 94.1%, this threshold minimizes false-positive results, which is particularly important in clinical contexts involving costly, invasive, or potentially harmful follow-up procedures, such as biopsies or advanced imaging. Although the corresponding sensitivity is relatively low (40.0%), this balance may be clinically acceptable for reducing overdiagnosis. Consequently, the 16.4 cut-off value may serve as a useful confirmatory tool following initial screening or as a decision aid within multi-step diagnostic algorithms in which clinical outcomes and specificity take precedence over sensitivity.

The promoter methylation analyses of tumor suppressor genes should be accompanied by gene expression studies to confirm the influence of hypermethylation of gene promoters on the production of tumor suppressor proteins. Suppressed expression of *CDO1* is expected if the promoter of a gene is hypermethylated. Promoter hypermethylation of the *CDO1* gene, accompanied by the suppressed expression of *CDO1* protein in immunohistochemistry, was demonstrated in patients with colorectal and gallbladder cancers [16, 22]. However, the *CDO1* promoter methylation level, accompanied by the *CDO1* gene expression in patients with GC or chronic gastritis, has not been investigated so far. Contrary to our expectation, we did not find a significant correlation between the *CDO1* promoter methylation level and immunopositivity score, neither in gastric tumors nor in biopsy samples of the chronic gastritis groups. The discrepancy in promoter methylation level of *CDO1* and *CDO1* immunopositivity score can be explained by possible different interpretations. The first is that different epigenetic modifications cooperate in the regulation of gene expression. Histone modifications such as acetylation and methylation are closely associated with DNA methylation in the control of gene expression. If histone modifications allow chromatin changes in the gene region, transcription may not be fully suppressed despite promoter methylation [23]. In this situation, the mRNA of the *CDO1* should be measured to understand whether the methylation over the promoter region of the *CDO1* affects its expression. The lack of *CDO1* mRNA analysis is a limitation of this study. The second is that if intracellular protein degradation is slowed down, the *CDO1* level may not change despite promoter methylation. The amount of intracellular cysteine is an important factor affecting the half-life of the *CDO1* protein [24]. At low cysteine levels, the ubiquitin-26S system can rapidly degrade *CDO1*. When the cysteine concentration is high, ubiquitination and subsequently degradation of *CDO1* are inhibited. The activity and expression level of *CDO1* are regulated depending on the intracellular cysteine concentration. Therefore, even if hypermethylation is observed in the *CDO1* gene, *CDO1* protein levels may be higher or lower than expected depending on the needs of the cell and the amount of intracellular cysteine [25]. As for the third comment, although the reference promoter, which is the region proximal to the transcription start site, is methylated, alternative promoter sites, even hundreds of base pairs away from the transcription start site, may sustain gene expressio [26, 27]. In addition to all these comments, tumor purity often causes erroneous findings. The samples collected from clinical settings are always a mixture of both cancer cells and non-cancerous stromal cells, immune cells, or epithelial cells. Methylation in cancer cells and non-cancerous cells is frequently different. Although the *CDO1* is methylated in tumor cells, non-cancerous cells may produce *CDO1.* In our opinion, methylation may trigger epigenetic, post-transcriptional, and post-translational modifications that affect the *CDO1* function; the intracellular degradation rate of the *CDO1* may have changed. If so, DNA repair slows down, mutations accumulate, and cancer progression accelerates. Immunohistochemical analysis shows the amount of protein, but it does not reflect its function. As another limitation of this study, the *CDO1* activity should be measured in addition to immunohistochemical analysis.

## Conclusion

The *CDO1* gene promoter levels are increased in GC patients and chronic gastritis patients with *H. pylori* infection. Promoter *CDO1* methylation, regardless of differences in protein expression, may play a significant role in the development of gastric cancer in HPP-chronic gastritis patients. Because of its reversible character, determination of DNA methylation changes in the promoter region of the *CDO1* gene may open a new window for early detection of GC, particularly in chronic gastritis patients with *H. pylori* infection. This is a preliminary study with a limited number of cases. A limitation of this study is the lack of a long follow-up period. We suggest examining the predictive and prognostic potential of the *CDO1* methylation as a non-invasive marker in chronic gastritis with *H. pylori* infection in large-scale prospective studies.

## Acknowledgements

We would like to thank to our Master’s Student Sıla Yağmur Beyaz, ‘Master’s Student, Istanbul University-Cerrahpasa, Cerrahpasa Medical Faculty, Department of Medical Biochemistry’, for her support of the manuscript preparation and Istanbul Yeni Yüzyıl University for supporting us in using their laboratory facilities.

## Supporting information

**S1 File. Ethic Approval English.**

**S2 File. Ethic Approval Turkish.**

**S3 File. Data Set**

